# Molecular Genetic Analysis of *RB1* gene among Sudanese children with Retinoblastoma

**DOI:** 10.1101/519306

**Authors:** Nada O. Ibrahim, Mahgoub Saleem, Entesar Eltayeb, Salwa Mekki, Elteleb G. Elnaim, Abeer Babiker Idris, Sanaa Bashir, Mohammed A. Salim, Mohamed A. Hassan

## Abstract

**BACKGROUND:** Retinoblastoma (RB), the commonest early childhood intraocular tumor, is most often related to mutations in the *RB1* gene with an incidence of 3% of all pediatric tumors. It has good prognosis if diagnosed early but it is life-threatening when diagnosed late.

**OBJECTIVE:** To study the Molecular Genetic Analysis of Retinoblastoma (RB) in Sudanese families.

**METHODS:** Thirty one (n=31) clinically and histopathologically diagnosed cases of RB attending Makkah Eye Complex (MEC) Orbit clinic (Khartoum, Sudan) were included in this Molecular Genetic RB Analysis. Fresh blood samples extracted from seven RB patients and 15 close families for DNA extraction and PCR were sent for Genetic Sequencing and *In silico* approach for “Exon 18 mutations” which is one of the highly mutated exons worldwide.

**RESULTS:** The majority of patients (41.9%) were below five years old. Females were 58.1%, males were 41.9%. Leukocoria was the commonest sign at presentation (41.9%). RB Unilaterality were in (77.4%) while Bilaterality in 19.4%. Both eyes were equally affected 50% each. The age at diagnosis time ranged from 0.02 to five years. Consanguinity of parents was very high (85.7%), the 1^st^ degree cousins were less (28.6%) while the 2^nd^ degree was high (57.1%). The patients’ ethnic background and geographical area were from seven different tribes; all belong to the Western Sudan. The molecular genetic study showed that exon 18 was free of mutation among the seven patients + their three relatives. The Functional Analysis and (SNPs) prediction study of exon 18 from NCBI data base showed that the various computational approaches used (*SIFT, PolyPhen-2, I-mutant* and *Project hope*) identified 16 reported mutations worldwide, three of which (rs137853292, rs375645171 and rs772068738) are major nsSNPs (non-synonymous) which might contribute to native *RB1* protein malfunction and ultimately causing carcinoma.

**CONCLUSION:** RB mainly affected children under five years and both sexes are equally affected. Unilaterality was predominant. Consanguinity plays a role in inheritance and the majority of patients were from Western Sudan. The most commonly detected deleterious mutations worldwide in exon 18 were not found in the Sudanese studies samples. Further screening for the highly reported mutations in exons 8, 10 and 14 or Next Generation Sequencing (NGS) are recommended. *In silico* tools are useful in studying the functional analysis of SNPs.

## Introduction

Retinoblastoma (RB) is a rare embryonic neoplasm of retinal origin and the most common intraocular tumour in children, with 3% incidence of all paediatric tumours and a frequency averaging 1:20.000 live born in different populations (1-9). It has good prognosis if diagnosed early but it is life-threatening when diagnosed late (1-5). The most common presenting sign is Leukocoria followed by strabismus (squint) (5, 7, 10).

The retinoblastoma gene (*RB1*) gene (MIM 180200) (3, 4) located on chromosome 13q14, consists of 27 exons (3, 4, 8, 11-14). It is a tumor suppressor gene and in its absence, chromosomal aberrations accumulate leading to tumor initiation, progression, and ultimately metastasis (6, 15-20). About one third of RB tumours are hereditary and bilateral, with a median age of one year at diagnosis. It is caused by an *RB1* constitutional mutation (M1) on one allele followed by a somatic *RB1* mutation, on the other allele (M2), leading to loss of function of the RB protein and initiation of tumor. This ‘two hit’ mechanism was first hypothesized by Knudson on 1971 (21-23). The other two-thirds of these tumours are unilateral, mostly non hereditary (86%) and have somatic inactivation of both *RB1* alleles with a median age of two years at diagnosis. The remaining 14% carry a germ line *RB1* mutation and are heritable retinoblastoma (2, 6, 7, 24). Patients with germ line *RB1* mutations can transmit the mutations to their offspring and are at increased risk for the development of secondary tumors (6, 8, 25-31).

The precise identification of the *RB1* mutations in each family with RB has been predicted to enhance the quality of clinical managemen**t** of the affected patient and relatives at risk (13, 32, 33). Children at risk undergo series of clinical examinations, including examination under anesthetic (EUA), so as to diagnose and treat tumors early. If *RB1* gene status of each family member is determined by molecular testing, only those relatives with the mutation require clinical surveillance, whereas non-carriers require no further examinations (25, 34). Thus molecular testing should be sensitive and economically feasible before it becomes routine clinical care (13, 35, 36). Genetic counseling and prenatal testing for pregnancies at risk are thus important (37, 38).

In Africa, many researches have been done regarding incidence, inheritance pattern, treatment outcomes and associated caners with retinoblastoma (39-45). Fewer researches have been conducted regarding mutational analysis of the *RB1* gene in African countries namely Morocco, Egypt, Algerian and Tunisian (46-50). Little research has been published in Sudan regarding retinoblastoma. Its incidence is estimated to be around 40-50 cases per year and it showed predilection distribution to Central and Western areas in Sudan (unpublished data by other colleges). Malik and El-Sheikh, 1979, studied 279 primary malignant tumors involving the eye and adnexa in Sudan, retinoblastoma formed 20.8% of it (around 58 cases) (51). Other studies showed that RB formed 2.7% and 4.8% out of the childhood cancers admitted to the institute of nuclear medicine in Gezira state (central Sudan) in a period of 5 years and 10 years respectively (52, 53). In fact, there is no study conducted in Sudan regarding the mutational analysis of the *RB1* gene in patient with retinoblastoma other than that conducted by Eltahir *et al,* 2011, where they showed contribution of Loss of heterozygosity (LOH) of the retinoblastoma gene (*RB1*) at two polymorphic intronic sites (intron 1 and 17) to cervical cancer (54). Therefore, the aim of this study is to screen for the most deleterious mutations reported world-wide in exon 18 (rs137853292, rs375645171 and rs772068738) in Sudanese families with positive Retinoblastoma member/s from the same ethnic background; taking into account that exon 18 is one of the highly mutated exons worldwide (1). To our knowledge, this is the first study done to screen exon 18 in Sudanese patients.

## Materials and Methods

### Study area

This study was carried out in Khartoum state at Makkah Eye Complex in Khartoum which is the largest hospital that provides eye care services for people from all parts of Sudan.

### Sampling

Out of all Sudanese patients diagnosed with RB (31 patients) attending Makkah Eye Complex during the year 2017, seven patients were selected randomly for genetic sequencing and analysis according to budget limitation. Three healthy family members with no retinoblastoma have been added as controls. Blood specimens were collected using EDTA-vacutainer tubes from the selected patients and controls. The specimens were preserved at −20 °C.

### Ethical considerations

Demographic data and the clinical information about each child and his family were collected through a close ended Questionnaire. A family pedigree was constructed for each family and was included in the questionnaire. All patients’ guardians were informed and consented to participate in the study before collecting the samples and were consented to publish the results of the study. Ethical approval was obtained from the ethical committee of Makkah Eye Complex.

### DNA extraction

For both patients and controls, DNA extraction was carried out using a kit from iNtRON Biotechnology. Genomic DNA extraction was confirmed by agarose gel electrophoresis and the DNA was kept at −20 °C until use.

### PCR amplification

Samples from seven patients and 15 relatives were subjected to amplification using the primer set in (Table 1) targeting *RB1* gene exon 18. Primers were synthesized and purchased from Macrogen Incorporation (Seoul, South Korea). Annealing temperature was adjusted using Maxime PCR PreMix Kit (i-Taq) 20 μl (INTRON Biotechnology, South Korea) on several runs of PCR. The adjusted temperatures are described in (Table 1). Amplification for the targeted region was done after addition of 14ul distilled water, 1ul DMSO, 3ul DNA sample and 1ul of each of the forward and reverse primers to the ready to-use master mix volume. PCR mixture was subjected to an initial denaturation step at 95 °C for 3 min, followed by 40 cycles of denaturation at 95 °C for 45 s, primer annealing at 55.6 °C for 60 s, followed by a step of elongation at 72 °C for 60 s, the final extension was at 72 °C for 7 min (55). PCR was repeated twice to yield enough DNA sample. The PCR products were checked and analyzed by 2% agarose gel electrophoresis at 100 V for 30 to 45 min and then bands were visualized by automated gel photo documentation system. *[Figure 1]* The seven patients and the three controls were subsequently selected for sequencing by the Sanger sequencing technique.

**[Table 1].** Primers used to amplify RB1 gene exon 18

| Primer's nucleotide sequence | PL* | AT* | Am* |
| --- | --- | --- | --- |
| **F:** 5' dTGACTTTATTTGGGTCATGTACCTT 3' | 25 | 55.6 | 360 |
| **R:** 5' dGCCACTGTCAATTGTGCCTA 3' | 20 | 55.6 |  |

PL\* Primer length in base pair, AT\* Annealing Temp, Am\* Amplicon size (bp)

**[Figure 1].**
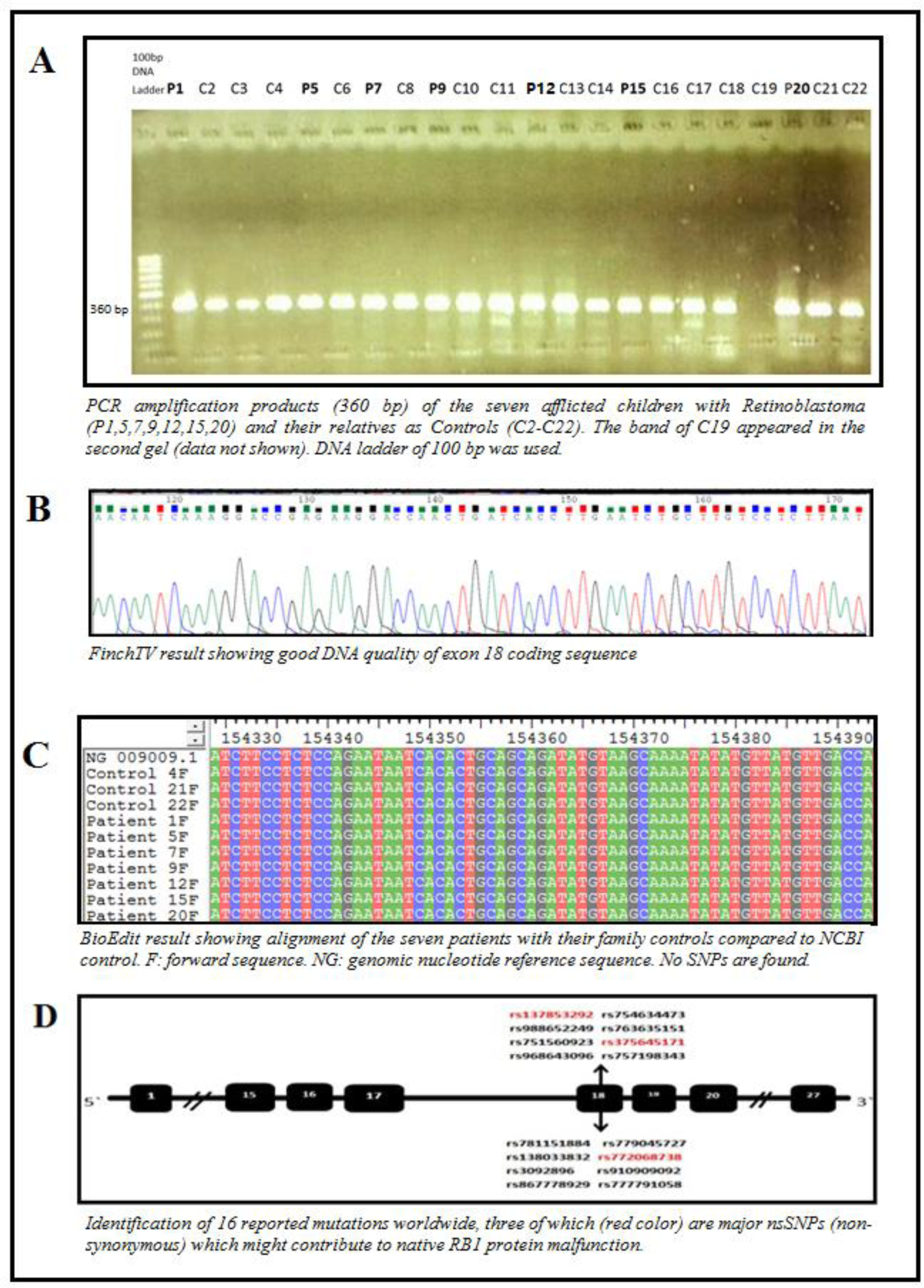
PCR and Bioinformatics results of the seven affected patients with RB and their control relatives. A: PCR. B: FinchTV. C: BioEdit.

### Sequencing of RB1 gene exon 18

Sanger sequencing was performed for the selected PCR products. Both DNA strands were sequenced by Macrogen Company (Seoul, South Korea).

### Bioinformatics analysis

FinchTV program version 1.4.0 (56) was used to view and check the quality of the two purified chromatogram (forward and reverse) nucleotide sequences from each sample. The NCBI Nucleotide database was searched for reference sequences. *RB1* nucleotide sequence (gene ref_ NG 009009.1) was obtained (57) and exon 18 was analyzed accordingly using nucleotide BLAST (Basic Local Alignment Search Tool) (58). BioEdit software (59) is used to find any apparent changes within the tested sequences through multiple sequence alignment.

## RESULTS

### Study population characteristics

#### Patient characteristics & clinical parameters

Thirty-one patients (n=31) diagnosed with Retinoblastoma (RB) attended Makkah eye complex (MEC) for the year 2017. The majority of patients (41.9%) when they attended the Orbit clinic and diagnosed with RB were below the age of 5 years (2-5 years), next were 1-2 years (29.0%) and only 3 patients (9.7%) were above the age of 5 years. *[Table 2, Figure 2]* Females slightly dominating males, 58.1% were females and 41.9% were males. *[Table 3, Figure 3]* Leukocoria seems to be the most common sign at presentation (n=13; 41.9%) despite incomplete records for all patients, while Phthisis, Enophthalmos and Retinal detachment were one case each. *[Table 4]* Unilaterality of the RB were described in (77.4%) while Bilaterality was less (19.4%); there was one missing record which will not affect the overall results. *[Table 5]* Right eyes and left eyes were equally affected 50% each. *[Table 5]* Regarding the seven RB cases, the demographic, characteristics and clinical features are summarized in *[Table 6 and 7].* Their age at diagnosis time ranged between seven days (0.02 year) to five years (mean age of 2 years ± 0.711 years). Again, the majority of patients (six patients) were below the age of five years, only one patient (14.3%) was about or above the age of five years. There were two males compared to five females. Five patients have unilateral sporadic tumor (i.e. no family history), one patient with unilateral familial tumor (his cousin is affected) and one patient with bilateral sporadic tumor (no family history at that time). The patient with the unilateral familial RB had a family history of a cousin who was diagnosed with an eye tumor at the age of seven months and died shortly three months later at the age of 10 months *[Table 6 and Figure 4].* Two patients have parents who are 1^st^ cousins, whereas four patients have parents who are 2^nd^ cousins. Only one patient his parents are not relatives. In summary: Consanguinity of parents was very high; found in six RB out of seven cases (85.7%), the 1^st^ degree cousins were less (28.6%) while the 2^nd^ degree was high (57.1%). The least was no Consanguinity of parents (14.3%). *[Figure 5]*

**[Table 2].** Age distribution at first RB diagnosis in MEC

| Age at diagnosis (years) | No. of patients | % |
| --- | --- | --- |
| 0-1 | 6 | 19.4 |
| >1-2 | 9 | 29 |
| >2-5 | 13 | 41.9 |
| >5 | 3 | 9.7 |
| Total | 31 | 100% |

Note: RB: Retinoblastoma; MEC: Makkah Eye Complex

**[Table 3].** Gender of RB patients attended MEC

| Sex | No. of patients | % |
| --- | --- | --- |
| M | 13 | 41.9 |
| F | 18 | 58.1 |
| Total | 31 | 100% |

**[Table 4].**
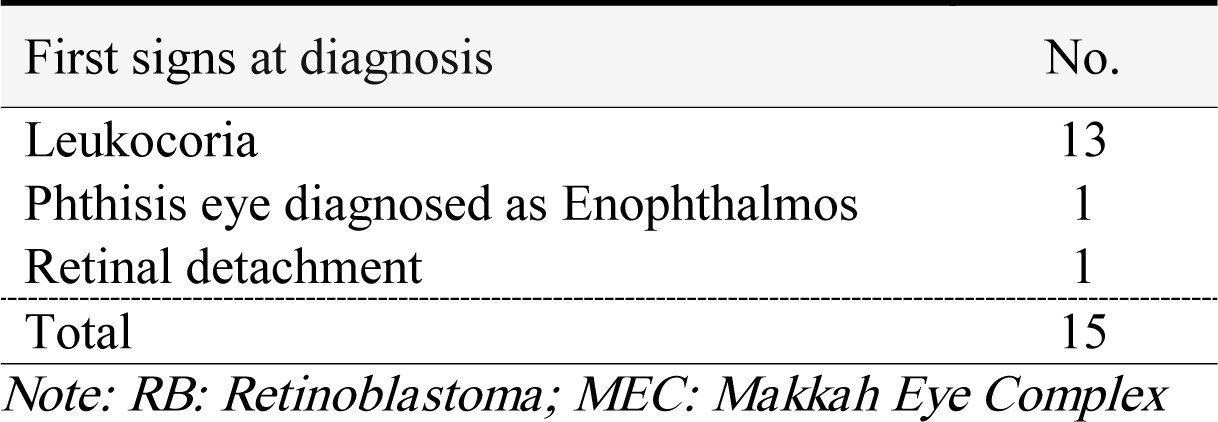
Distribution of RB First sign at diagnosis in MEC

**[Table 5].** Laterality Distribution of RB in MEC

| Laterality | No. | % | OD | % | OS | % |
| --- | --- | --- | --- | --- | --- | --- |
| Unilateral | 24 | 77.4 | 12 | 50 | 12 | 50 |
| Bilateral | 6 | 19.4 | - | - | - | - |
| Total | 30 | 96.8% |  |  |  |  |

Note: RB: Retinoblastoma; MEC: Makkah Eye Complex; OD: Oculus Dextrus: right eye; OS: Oculus Sinister: left eye

**[Table 6].**
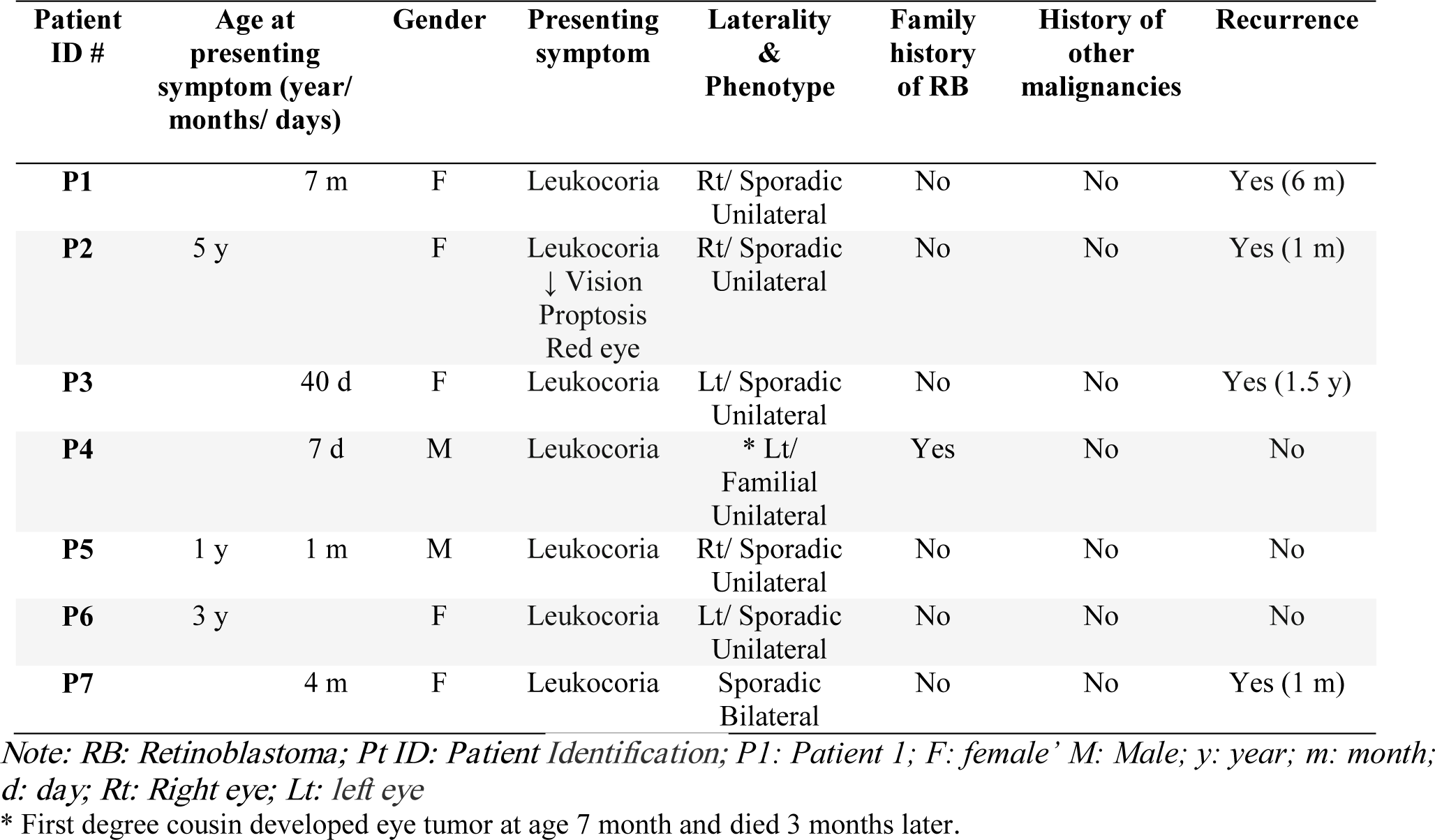
RB patients’ demographic and clinical features for the 7 studies cases

**[Table 7].** The demographic, characteristics and clinical features of the seven RB patients attended MEC (2017)

| Variable |  | Frequency (%) |
| --- | --- | --- |
| Onset | ≤1 year | 4 (57.1%) |
|  | 1-2 years | 1 (14.3%) |
|  | 2-5 years | 1 (14.3%) |
|  | >5 | 1 (14.3%) |
| Sex/Gender | Males | 2 (28.6%) |
|  | Females | 5 (71.4%) |
| Family history | Ocular cancer | 1 (14.3%) |
|  | Other cancer | 0 (0%) |
| Consanguinity of parents | 1<sup>st</sup> degree cousins | 2 (28.6%) |
|  | 2<sup>nd</sup> degree cousins | 4 (57.1%) |
|  | None | 1 (14.3%) |
| Tribe | Baramka | 1 (14.3%) |
|  | Rezaigat (Kurdofan/Darfor) | 1 (14.3%) |
|  | Jammoeyya (Kurdofan) | 1 (14.3%) |
|  | Zaghawa (Darfor) | 1 (14.3%) |
|  | Fallata (Darfor) | 1 (14.3%) |
|  | Mesaireya (Kurdofan/Darfor) | 1 (14.3%) |
|  | Kawahla | 1 (14.3%) |
| Geographical region | Central Sudan<sup>[1]</sup> | 0 |
|  | Western Sudan | 7 (100%) |
|  | Northern Sudan | 0 |
|  | Eastern Sudan | 0 |
| Tumor site | Unilateral | 6 (85.7%) |
|  | Bilateral | 1 (14.3%) |

<sup>[1]</sup> Comprising both Khartoum and Al Gezira

**[Figure 2].**
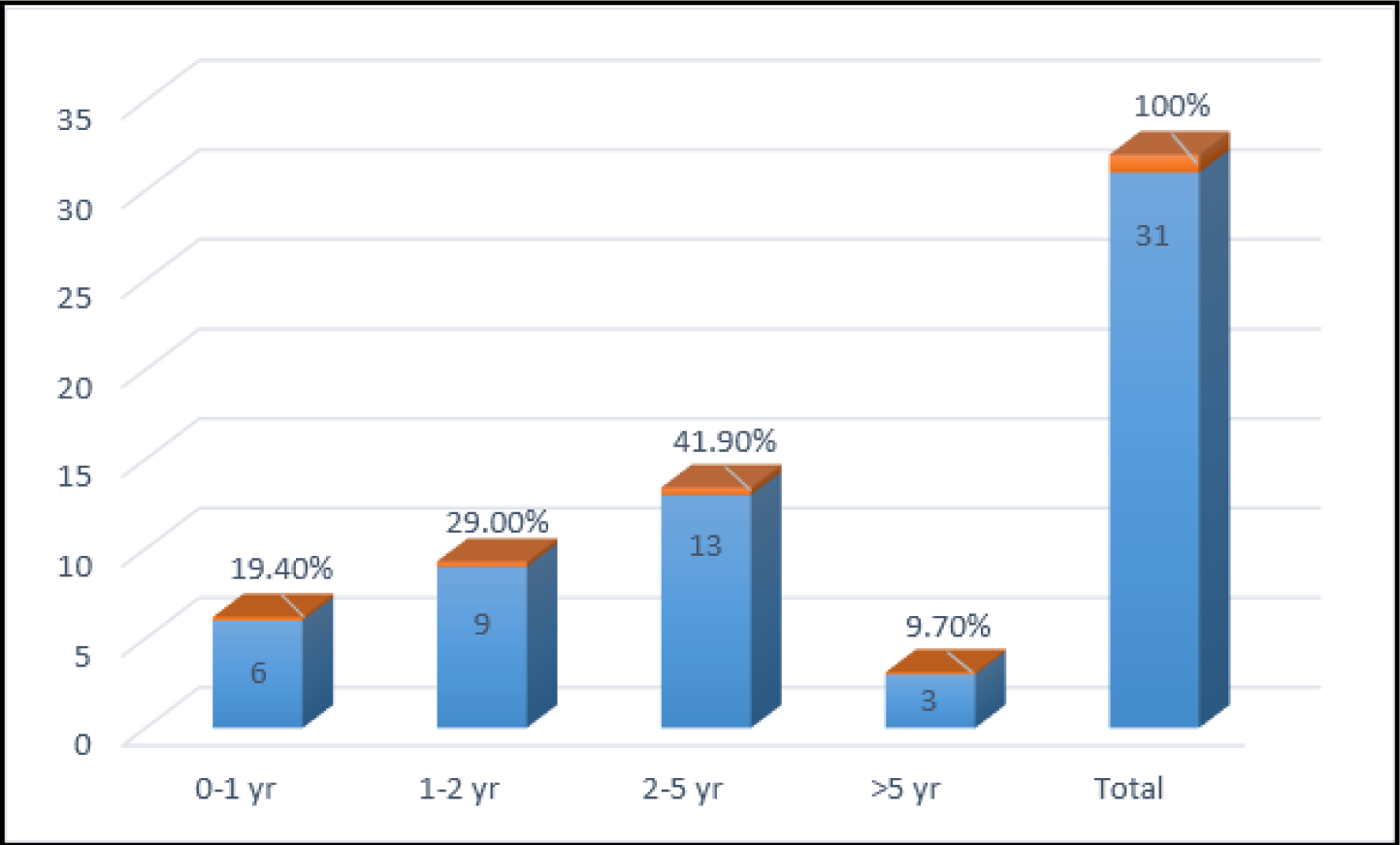
Age distribution at first RB diagnosis in MEC Note: RB: Retinoblastoma; MEC: Makkah Eye Complex

**[Figure 3].**
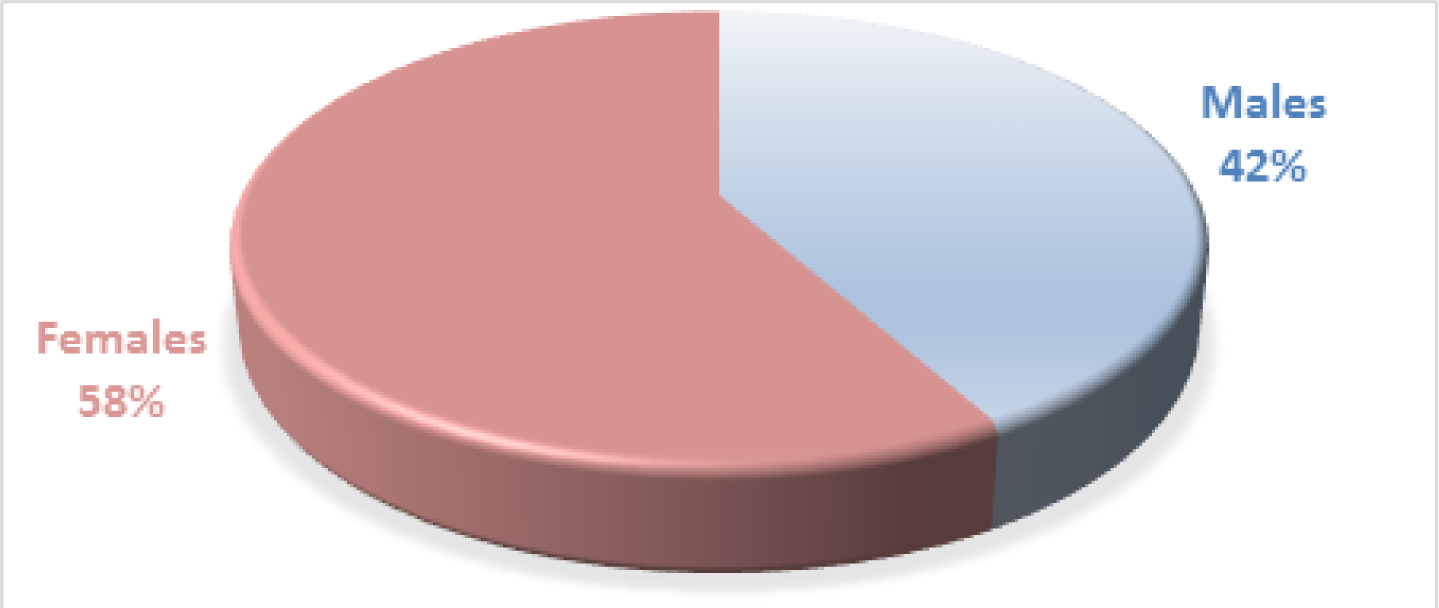
Gender distribution of Retinoblastoma in MEC Note: MEC: Makkah Eye Complex

**[Figure 4].**
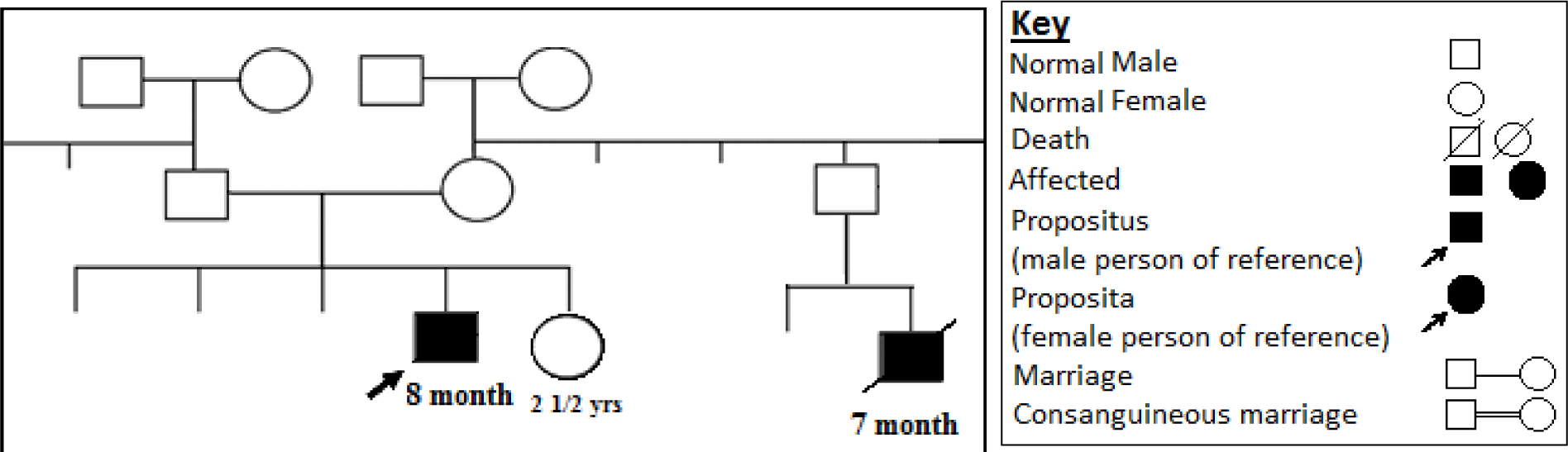
*Pedigree of an 8 month old RB* patient with a unilateral familial RB who has a cousin that is diagnosed with an eye tumor at the age of seven months and died shortly after three months at 10 months of age.

**[Figure 5].**
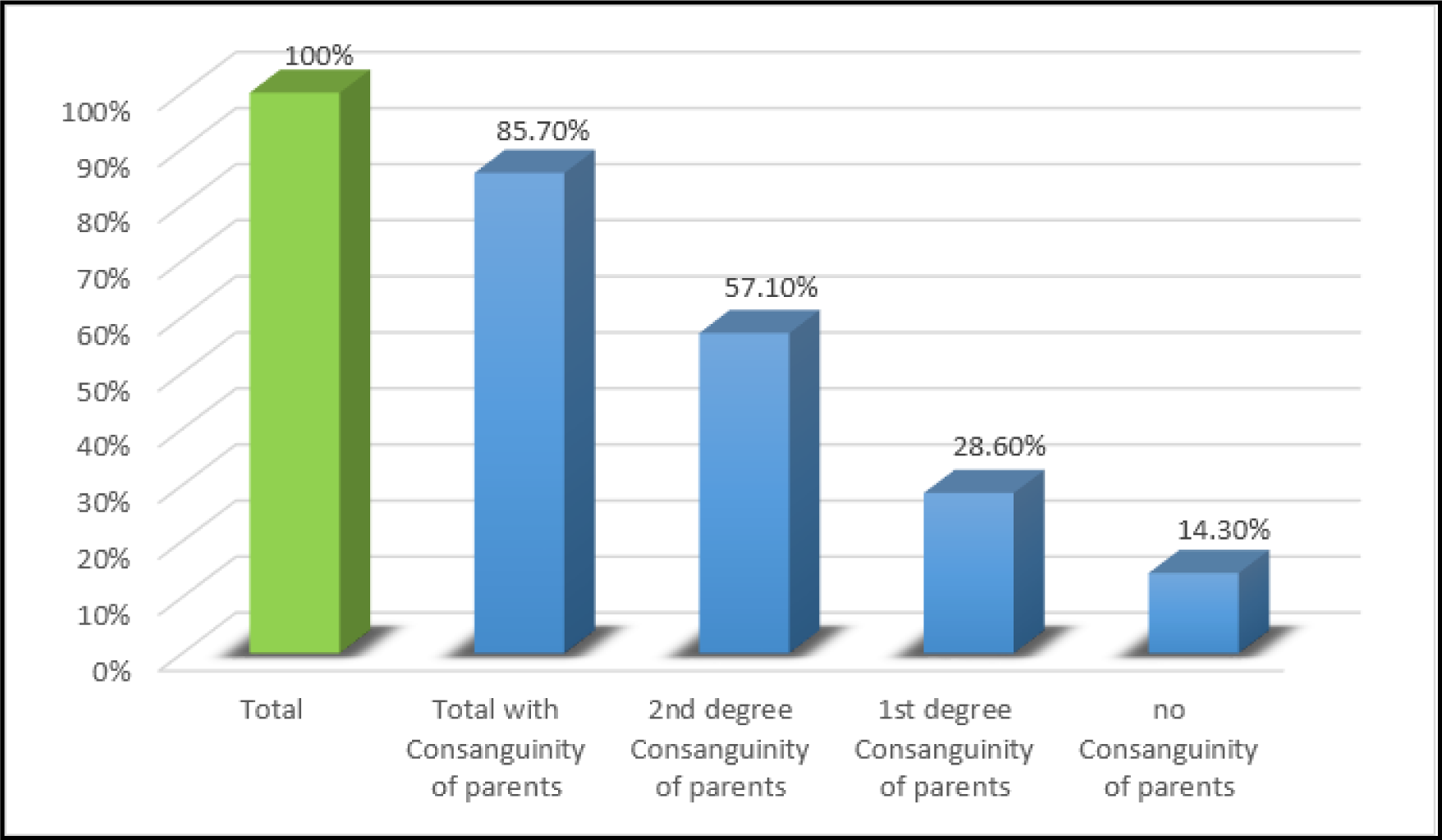
Distribution of Consanguinity of Parents in Retinoblastoma cases of MEC Note: MEC: Makkah Eye Complex

Regarding laterality of the disease, our small sample again showed that unilateral RB (85.7%) is highly dominant to bilateral disease (14.3%). No distant metastasis was reported in our samples but there were four recurrences out the seven samples (57.14%).

Regarding ethnic background and geographical area, patients were from seven different tribes; but all of them (100%) belong to the Western Sudan [Baramka (Kurdofan/Darfor), Rezaigat (Kurdofan/Darfor), Jammoeyya (Central/Kurdofan), Mesaireya (Kurdofan), Kawahla (Kurdofan), Zaghawa (Darfor) and Fallata (Darfor and Western Afric). *[Table 7]* Geographical area is not definite as most of these tribes are nomadic with known habits to move across the country with their animals. *[Table 7]*

### PCR products of exon 18

The PCR products (360 nucleotide long) of the seven afflicted children with Retinoblastoma (P1, 5, 7, 9, 12, 15, 20) and their relatives as Controls (C2-C22) are shown in figure 1A. All 22 samples yielded sufficient quality bands (sample # 19 did not show a PCR product in this gel picture but gave a clear band in another run. Picture is not shown).

### Bioinformatics result analysis

The sequencing data was checked for consistency and quality by FinchTV as shown in figure 1B. Our result showed consistency and good quality for all 10 sequences. By using the multiple sequence alignment tool BioEdit, the analysis of seven tested patients and three family controls compared to NCBI reference sequence RefSeq (NG 009009.1) revealed no nucleotide change as shown in figure 1C.

## DISCUSSION

In our study, the number of RB patients attending Makkah Eye complex in the year 2017 was 31. The majority of our patients were below the age of five years and there is slight predilection for the female gender with a male to female ratio of 1:1.4. Despite the incomplete records, Leukocoria seems to be the most common presenting sign at presentation (13 patients). This finding is consistent with the result reported by K Shahraki *et al.* in Iran that the Leukocoria represent the major complaints of RB patients (60). Regarding laterality of the disease, our results were slightly different than that in the literatures where unilateral tumors occupied 77.4% and bilateral tumors 19.4% compared to ∼ 60% unilateral and 40% bilateral worldwide (2, 6, 7, 24). For unilateral tumors, right and left eyes are equally affected and seems to have no predilection to one side. Only seven patients attended for treatment and follow-up were selected for our molecular genetic study due to budget limitation. Further screening for the highly reported mutations in exons 8, 10 and 14 or NGS (Exom sequencing) are recommend with larger sample size or a whole family (including the child, parents & siblings).

Most of our patients had a unilateral sporadic disease (five out of seven). Familial history of an eye tumor was found in one case with a unilateral RB and parents who are 2^nd^ degree cousins. The patient’s relative, who was a cousin, had an eye tumor at the age of seven month and died shortly at the age of 10 months old (figure 4). This supports the literature in that familial RB can presents as unilateral disease (2, 6, 7, 24). The other patient (two years old) with bilateral sporadic tumor and whose parents are not relatives had a younger sibling who was one month old at the time of study and thus no family history was demonstrated at that time. This family needs follow up for possible sibling affection with RB. Regarding consanguinity, our results showed that RB can occur in 2^nd^ degree relatives as well as in non-relatives. This demonstrates that RB has no predilection to close relationship per say, it can occur in any family. Regarding ethnic background and geographical area, all seven studied families were from Western Sudan. Although this is a small sample number, it is comparable with a study performed by a college (unpublished) which demonstrated that RB is present in Western Sudan (Kordofan and Darfour) in a slightly higher percentage (40.6%) than Central Sudan (Khartoum and Al Gezira) (34.4%). Again for unilateral tumors, right and left eyes were equally affected and seems to have no predilection to one side. Recurrences occurred in four patients either immediately after six months of chemotherapy or later, indicating late presentation and spread of the disease or ineffectiveness of the chemotherapy.

Mutations in exon 18 is considered to be one of the hotspots reported world-wide in a searchable database (1, 61-65). The various computational approaches used (SIFT, PolyPhen-2, I-mutant and Project hope) identified 16 reported mutations worldwide, three of which (rs137853292, rs375645171 and rs772068738) are major nsSNPs (non-synonymous) which might contribute to native *RB1* protein malfunction and ultimately causing carcinoma (Figure 1D). In this study, these targeted mutations were not found in exon 18 of *RB1* gene of our Sudanese samples. However, mutations in *RB1* gene are random and could be found in other exons as well due to heterogenicity of the disease, and thus screening of all exons is more appropriate. In addition, several previous studies from different countries are in agreement with our finding and they revealed many mutations across *RB1* gene but not in exon 18 in patients with retinoblastoma (unilateral, bilateral, trilateral, sporadic and/or familial RB cases) (66-68).

To our knowledge, there is no study conducted in Sudan regarding the mutational analysis of the *RB1* gene in patient with retinoblastoma other than that conducted by Eltahir *et al,* 2011, where they showed contribution of Loss of heterozygosity (LOH) of the retinoblastoma gene (*RB1*) at two polymorphic intronic sites (intron 1 and 17) to cervical cancer (54) and thus, this is the first study done to screen exon 18 of *RB1* gene in Sudanese patients.

## CONCLUSION

RB mainly affected children under five years and both sexes are equally affected. Unilaterality was predominant. Consanguinity plays a role in inheritance and the majority of patients were from Western Sudan. The most commonly detected mutations worldwide in exon 18 were not found in our Sudanese studies samples. Further screening for the highly reported mutations in exons 8, 10 and 14 or NGS (Exom sequencing) are recommended with larger sample size or a whole family (including the child, parents & siblings). In silico tools are useful in studying the functional analysis of SNPs. To our knowledge, this is the first study performed to screen exon 18 in Sudanese patients.

